# Rapid gain and loss of a chromosome drives key morphology and virulence phenotypes in *Histoplasma*, a fungal pathogen of humans

**DOI:** 10.1101/2025.05.23.655702

**Authors:** Sarah Heater, Mark Voorhies, Rosa A. Rodriguez, Bevin C. English, Anita Sil

## Abstract

Heritable phenotypic switches are fundamental to the ability of cells to respond to specific conditions. Such switches are key to the success of environmental pathogens, which encounter disparate conditions as they transition between the environment and host. We determine that the copy number of chromosome seven in the thermally dimorphic fungus *Histoplasma* dramatically affects the rate of transition. Though *Histoplasma* is haploid, a second copy of this chromosome is present in natural isolates of multiple *Histoplasma* species and is gained and lost at a high rate. Cells carrying two copies of this chromosome exhibit aspects of the environmental transcriptome even under host-like conditions and have a competitive advantage in the transition to the environmental form. Conversely, these cells are considerably less virulent than euploid cells and have a competitive disadvantage in the mouse model of infection. Chromosome seven contains a previously unstudied transcription factor that, when overexpressed in euploid *Histoplasma*, is sufficient to promote some of the key phenotypes of aneuploidy. We hypothesize that rapid gain and loss of this chromosome benefits *Histoplasma* by increasing phenotypic variation, thus helping populations of cells survive abrupt transitions between environment and host.

## Introduction

Genetic diversity within a population can enable survival, since a portion of the population may survive conditions lethal to most of the community. Genome instability is one mechanism by which a community can rapidly diversify, preparing it to survive subsequent fluctuations. One form of genome instability is copy number variation, ranging from changes in copy number of the entire genome, a particular chromosome, or a specific gene. Copy number variation has been associated with improving fungal fitness in a wide range of stress conditions, including exposure to antifungals, high temperature, and mammalian infection^1–6^.

Exposure to extreme fluctuations is fundamental to the lifecycle of environmental pathogens such as *Histoplasma* species, which transition between the environment and the mammalian host. *Histoplasma* grows as sporogenous hyphae in soil at environmental temperatures. When aerosolized and inhaled by a human or other mammal, *Histoplasma* transitions to growth as pathogenic yeast in response to body temperature. Upon return to the environment, potentially via excretion in guano^7^ or after host death^8^, *Histoplasma* transitions back to its hyphal form. In vitro, simply shifting the temperature from ambient room temperature to body temperature is sufficient to trigger this morphology transition. Understanding the molecular basis of morphologic transitions in response to temperature is fundamental to understanding *Histoplasma* pathogenesis.

*Histoplasma* is considered a high priority pathogen by the WHO and is found globally, with specific regions of hyperendemicity^9,10^ where most of the human population may be exposed to *Histoplasma* within their lifetimes^11,12^. A recent analysis suggested that *Histoplasma* incidence among people with AIDS in Latin America may be similar to that of tuberculosis^13^. Despite this high prevalence, infections are often misdiagnosed and it is considered a neglected disease^13^.

Recently, some progress has been made in our understanding of the molecular mechanisms driving *Histoplasma* towards the pathogenic yeast or the environmental hyphal morphology in response to temperature changes, including the discovery of several transcription factors (TFs) that drive each morphology^14–19^. However, one difficulty in these investigations has been a lack of consistency in these transitions in vitro. While temperature is sufficient to trigger transitions between yeast and hyphal growth in the laboratory, the rate at which this transition occurs has not been fully consistent for unknown reasons. Differences in the rate of transition were previously misattributed to the presence or absence of the gene *MSB2*^20,21^.

Enabled by our recent whole genome assembly^22^, we found that frequent gain and loss of a specific chromosome in this haploid organism affects the rate of hyphal formation in response to temperature shift. Aneuploid *Histoplasma*, carrying an additional copy of chromosome 7, is biased towards growth as hyphae and rapidly transitions from yeast to hyphae in response to ambient temperature. Conversely, euploid *Histoplasma* is biased towards growth as yeast and undergoes a much slower transition to hyphae in response to temperature. We discovered that copy number variation of this chromosome is present in many natural and laboratory isolates, and that this aneuploidy is gained and lost at a high rate. Cells carrying two copies of chromosome 7 are significantly less virulent in the mouse model of infection than cells carrying a single copy. A transcription factor on chromosome 7 is sufficient to drive some of the phenotypes associated with an extra copy of chromosome 7, including transcriptional upregulation of a conserved set of genes associated with the hyphal morphology. This work sets the stage for understanding how variation within a population affects the ability of this important fungal pathogen of humans to thrive within the environment and host.

## Results

### *Histoplasma* morphology correlates with gain and loss of chromosome 7

Morphological shifts are critical to the pathogenic lifecycle of *Histoplasma*. As such, morphological bias—i.e., the phenotypic trait of favoring a particular morphology—is of great potential natural importance. We found that some *Histoplasma ohiense* strains were yeast-biased, continuing to grow as yeast for weeks in hyphae-inducing conditions, while other strains were more hyphal-biased, converting rapidly from yeast to hyphal growth when transitioned to ambient temperature. This morphological bias could be clearly observed by microscopy of cells grown in liquid culture (Fig. 1A) as well as colony morphology (Fig. 1B). Previously, a similar yeast-biased phenotype in liquid culture was mistakenly attributed to mutation of the *MSB2* gene, a finding that we could not replicate^20,21^. We thus subjected yeast- and hyphal-biased strains to whole genome sequencing.

**Figure 1:**
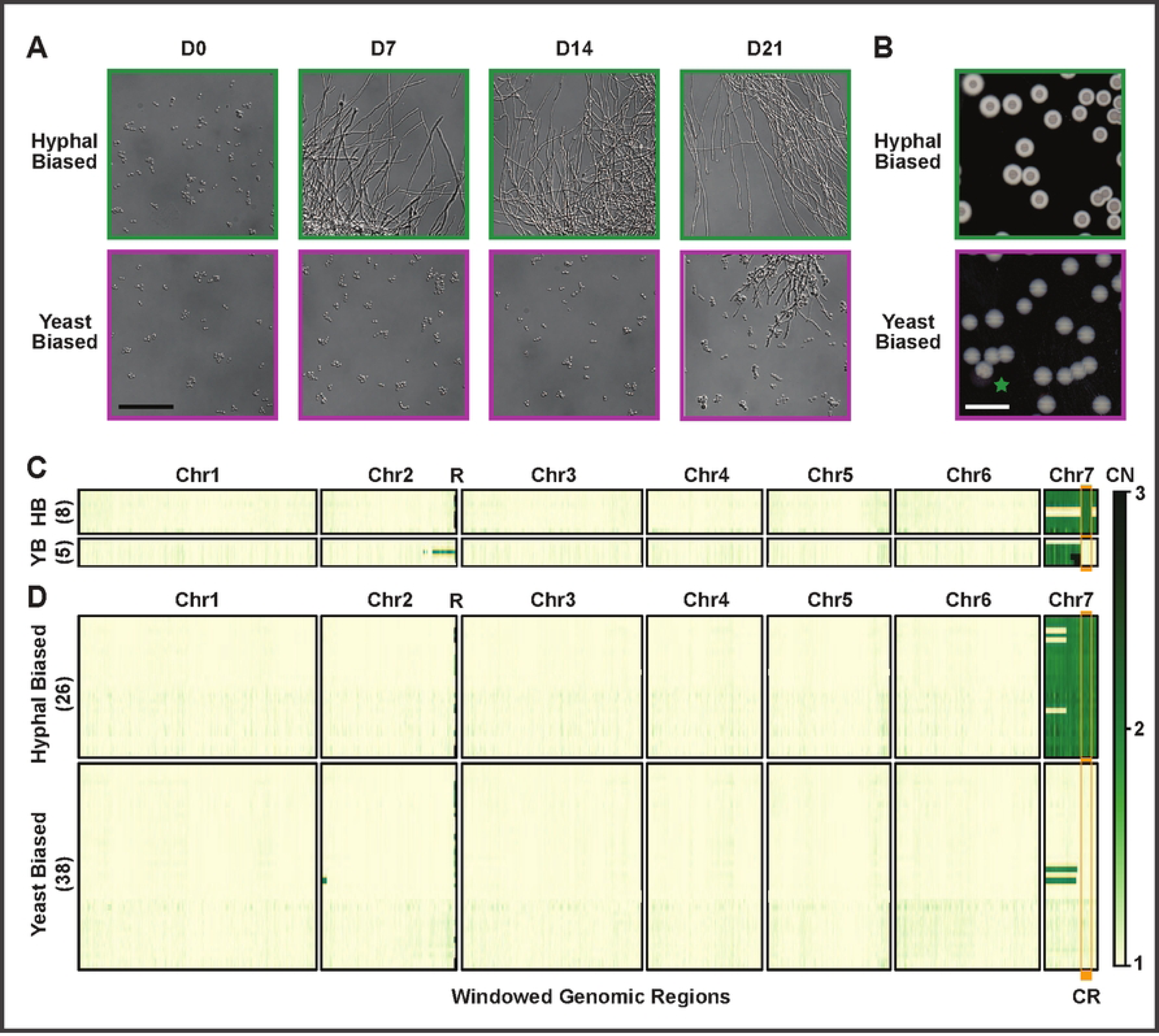
*Histoplasma* morphology correlates with aneuploidy of chromosome 7. Morphological bias of *Histoplasma* strains can be observed both microscopically as cell morphology and macroscopically as colony morphology. A) Representative microscopy images from strains with each morphology bias over a 21-day shift from 37°C to 25°C. Bar indicates 50 µm. B) Representative images of colony morphology, including one yeast-biased colony with a potential sector of hyphal bias (green star). White bar indicates 10 mm. Green and magenta are used in A and B to indicate morphology bias as well as throughout this study. C-D) Windowed copy number (CN) based on normalized sequencing coverage of strains, excluding LTR retrotransposon blocks. Strains were categorized as hyphal-biased (HB) or yeast-biased (YB) prior to Illumina sequencing. The number of strains in each category is listed in parentheses on the y axis. A vertical orange bar is used to indicate the critical region (“CR”), the region of Chr7 that best correlates with morphology bias. The letter R indicates the location of ribosomal DNA on Chr2. C) Strains used to define the critical region and control strains. These include 6 strains with a partial second copy of Chr7. Closely related control strains with a full single copy or full double copy of Chr7 are also shown. D) Strains generated alongside one or more sibling strain of the opposite morphology bias. Correspondence between morphology bias and CR copy number is greater than expected by chance, as is correspondence between morphology bias and Chr7 copy number among strains with full 1X or full 2X Chr7 coverage. Fisher exact p-value < 1×10^-7^ for both comparisons. All strains used here are derived from *H. ohiense* clinical isolate G217B. Additional information about the sources of strains in C and D is shown in Fig. S1.

Based on 6 strains with a partial second copy of chromosome 7 (Chr7), we defined a 334 kb region of Chr7, which we named the critical region, that correlated with hyphal bias (Fig. 1C). Yeast-biased strains universally lacked an extra copy of this region. Hyphal biased isolates universally contained a second copy of this region.

We observed that yeast-biased isolates could spontaneously give rise to hyphal-biased isolates, and vice versa. Sixty-four strains, each generated concurrently with matched sibling strain(s) of the opposite morphology bias, were derived in multiple ways and from multiple genetic backgrounds (Fig. S1). To ensure a robust assessment of the correlation between morphology and copy number variation (CNV), some strains were selected randomly and subsequently assessed for morphology bias while other strains were selected based on morphology bias. Strains were then subjected to whole genome sequencing. Hyphal bias generally correlated with a second copy of Chr7. The association between duplication status of the Chr7 critical region and morphology bias was 100% (Fig. 1D).

### Chromosome 7 aneuploidy is present in multiple *Histoplasma* isolates and species

Given the abundance of observed instances of Chr7 aneuploidy and the perfect correlation between CNV and morphological bias observed in the laboratory, we hypothesized that this aneuploidy would be relevant in natural populations. From analysis of previously published genome sequences^23–25^, we found duplication of the full chromosome or the critical region in 16% of *Histoplasma* natural isolates of multiple species (Fig. 2A, S2A-C). These data included CNV in the 6^th^ largest chromosome of *H. mississippiense*, which is syntenic to *H. ohiense* Chr7^22^. Individual natural isolates were subjected to short-term culturing, and then progeny were assessed for morphology bias and aneuploidy. Progeny displayed rough or smooth colony morphology, generally based on presence or absence of a hyphal ring indicating hyphal bias (Fig. 2B). Sequencing of these isolates revealed that for 14/16 sequenced strains, colony morphology bias correlated with the CNV of interest, for both *H. mississipiense* and *H. ohiense* (Fig. 2C-D). The remaining two sequenced strains displayed a unique phenotype of increased colony size and a rough variegated colony surface appearance rather than a rough fuzzy hyphal ring around the colony (4-Rough, Fig. 2B). These two strains had a small CNV on chromosome 5 (Fig. 2D). While CNV was observed among natural isolates in regions other than the aneuploid chromosome, CNV was by far most common in the aneuploid chromosome and was especially common in *H. ohiense* in the critical region (Fig. S2A-C).

**Figure 2:**
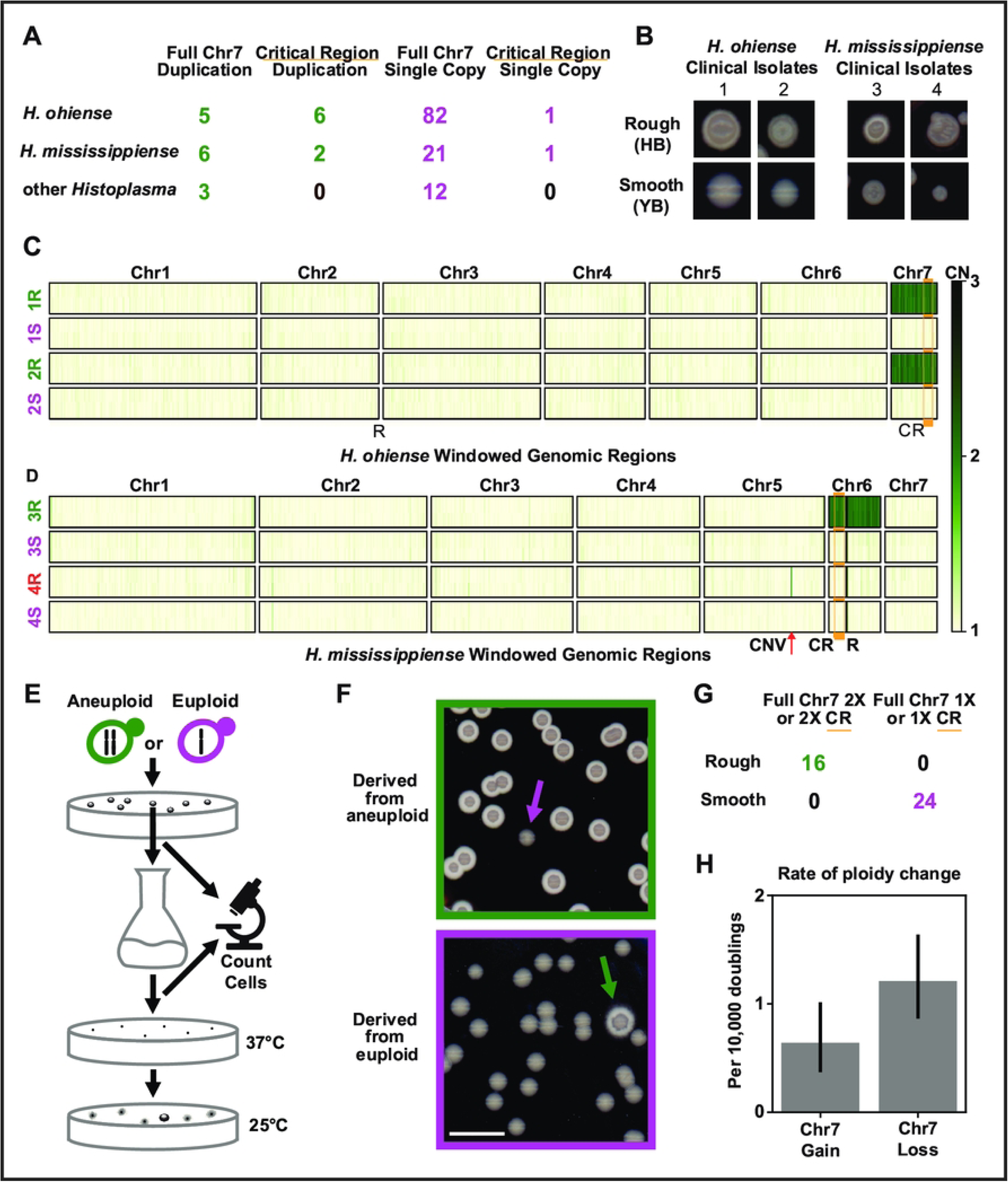
Ploidy variation correlating with morphology occurs in multiple *Histoplasma* species and is rapidly gained and lost. A) The Chr7 aneuploidy is present in natural population isolates of *H. ohiense*, *H. mississippiense*, and other *Histoplasma* sequenced by multiple previous groups. Columns from left to right indicate the number of isolates with a full second copy of Chr7, a second copy of a region of Chr7 including the critical region, only a single copy of Chr7, and a second copy of a region of Chr7 not including the critical region. B) Distinct colony morphologies were identified among progeny of two additional *H. ohiense* natural isolates (#1: CI_4, #2: CI_9) and two *H. mississippiense* natural isolates (#3: CI_43, #4: UCLA). For each of these four parental groups, morphology bias was categorized as smooth or yeast-biased (YB, e.g. category 1S from isolate 1) and rough or hyphal-biased (HB, e.g. category 1R). Progeny were subsequently sequenced, and copy number through the genome of these isolates is shown in for *H. ohiense* strains (C) and *H. mississippiense* strains (D). Strain categorization names are colored based on ploidy of strains in this category, with strains having a small CNV on chromosome 5 noted in red. The location of this small CNV is noted by “CNV” and a red arrow. As in Fig. 1, critical region (CR) is noted by orange bar and rDNA is noted by the letter R. E) Schematic of how cells were briefly grown up (over an average of 31 doublings) while doublings were counted. Morphology bias among progeny was tested to determine Chr7 copy number gain-and-loss rates. F) Example images of colonies from gain-and-loss rate experiments as described in E. Cells derived from an aneuploid parent (above, green) and colonies derived from a euploid parent (below, magenta) each showing one progeny colony (indicated by an arrow) that converted in morphology bias. Bar indicates 10 mm. Conversion in morphology bias was used as a proxy for converting in ploidy. G) Forty colonies from the assays used to determine gain and loss rates were sequenced to confirm morphology-CNV correlation. This sequencing is also included in Figure 1 and S1. H) Rates of Chr7 gain and loss determined by cell doublings and rate of morphology bias switching. Error bars show 95% confidence interval.

### The chromosome 7 aneuploidy is rapidly gained and lost

To assess the rate of gain and loss of the Chr7 aneuploidy, aneuploid and euploid parental cells were subjected to short-term culturing and ploidy of their progeny was assessed (Fig. 2E). Importantly for this analysis, aneuploid and euploid yeast had indistinguishable growth rates (Fig. S3A). We used colony morphology as a proxy for Chr7 aneuploidy and validated this assumption by sequencing 40 progeny colonies and obtaining a perfect correlation between ploidy of the critical region and morphology bias (Fig. 2F-G, S3B). We found that the rates of gain and loss of the Chr7 aneuploidy were 5.0×10^-5^ and 1.1×10^-4^ per doubling, respectively (Fig. 2H). Surprisingly, these rates are significantly higher than the estimated rate of chromosome gain in haploid *Saccharomyces* and are comparable to the rate at which diploid *Saccharomyces* develops any individual aneuploidy^26–28^.

### Chromosome 7 aneuploidy confers a competitive advantage during the yeast-to-hyphal shift and a disadvantage during the hyphal-to-yeast shift

Given the high prevalence of the Chr7 aneuploidy and the high rate of gain and loss, we went on to assess if there might be conditions in which the aneuploidy confers a competitive advantage in a mixed population. We co-cultured an equal number of euploid and aneuploid cells and determined relative abundance of each population during temperature shift. A morphology score was assigned to each timepoint as previously described^22^. We found that in pooled competitions, the hyphal-biased aneuploid strains took over the population during the shift from yeast to hyphae at the timepoints at which yeast were observed to convert to hyphae in these mixed ploidy cultures (Fig. 3A-C). Conversely, it was the yeast-biased euploid strains that took over the population during the reverse shift from hyphae to yeast (Fig. 3D-F). In both cases, this increased abundance could be due primarily to the morphology bias of each strain.

**Figure 3:**
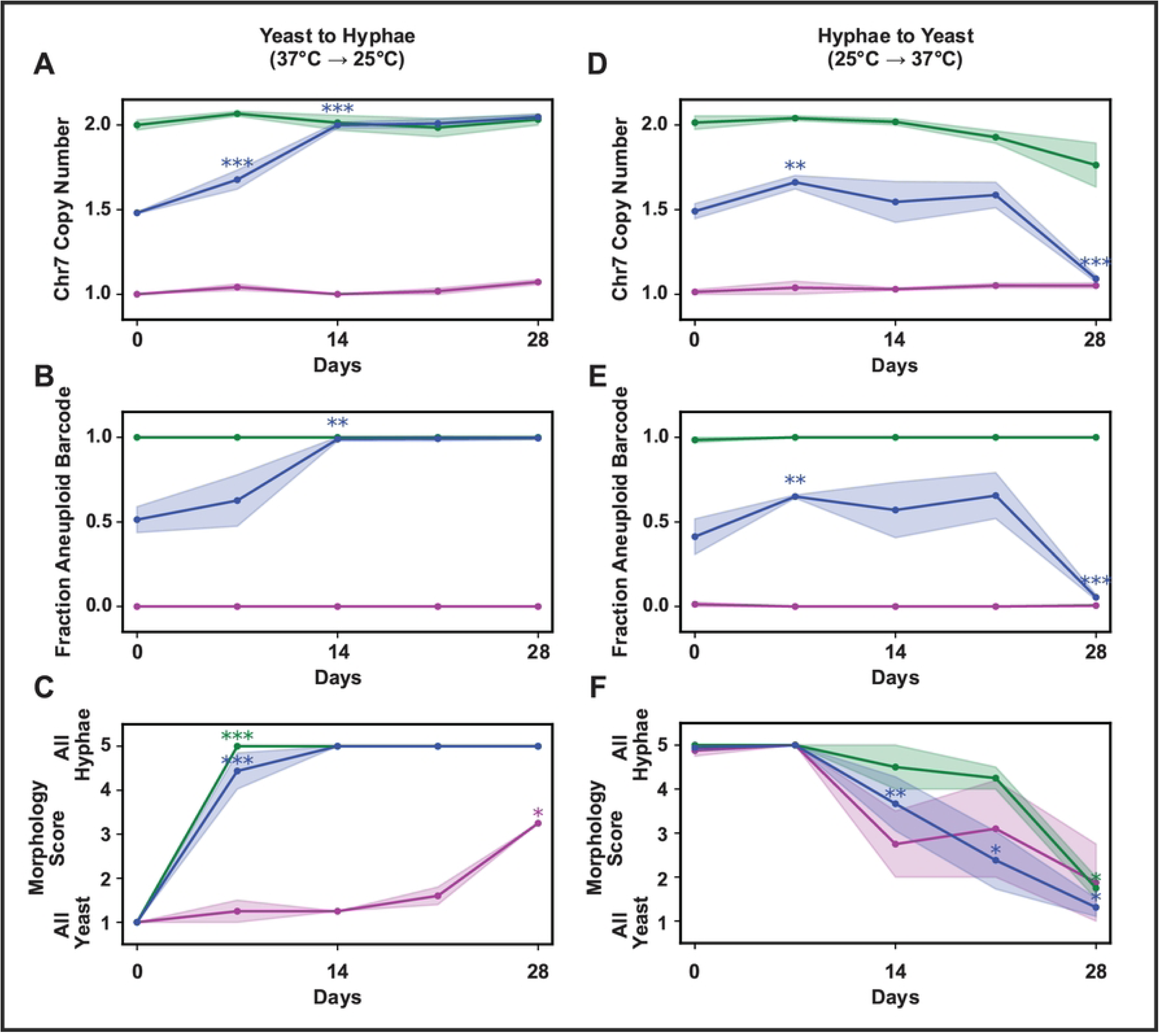
Chr7 aneuploidy confers a competitive advantage during the shift to the environmental form but confers a disadvantage during the shift to the host form. Over the course of four weeks, yeast was transitioned to hyphae (moved on day 0 from yeast inducing conditions at 37 °C with added CO_2_ to hyphal inducing conditions at 25 °C without added CO_2_) and hyphae were transitioned to yeast (moved on day 0 from hyphae inducing conditions at 25 °C without added CO_2_ to yeast inducing conditions at 37 °C with added CO_2_). Starting populations for these transitions included 50%-50% mix of aneuploid and euploid cells (blue), as well as pure aneuploid (green) and pure euploid (magenta) populations. A-C show the yeast-to-hyphae transition while D-F show the hyphae-to-yeast transition. A&D) Chr7 copy number throughout morphological transitions based on normalized sequencing coverage. B&E) Population ratios as fraction of aneuploid starting strain from SNP-based barcoding throughout transitions. C&F) Morphology, assessed by microscopy followed by image categorization, throughout morphological transitions. Morphology score ranged from 1 to 5 corresponding to all yeast (1), majority yeast (2), mixed (3), majority hyphae (4), all hyphae (5). We note as a caveat that cultures throughout the later portion of the hyphae-to-yeast transition exhibited notable flocculation which may increase morphology score variability. For each subfigure, asterisks show significance by t-test of change versus the prior point, colored based on population.

We also determined that significant Chr7 gain or loss within the population did not explain the observed changes in Chr7 copy number in these experiments. We used two spontaneously occurring non-coding SNPs as “barcodes” to track the starting strains. Euploid and aneuploid counterparts were generated for strains differing by only these barcodes and the appropriate pairs were mixed and subjected to morphology analysis and sequencing. Change in barcode ratios corresponded with change in Chr7 copy number, indicating that changes in the relative ratio of euploid to aneuploid cells was due to outcompetition rather than widespread Chr7 gain or loss over the course of the experiment (Fig. 3A-B,D-E).

In addition to mixed cultures, we also examined pure aneuploid and pure euploid cells under both transitions. In general, these cells maintained their ploidy state throughout the morphological transitions. However, we observed minor conversion of aneuploid to euploid cells in the hyphae-to-yeast transition as observed by the decrease in Chr7 copy number in the aneuploid population (Fig. 3D). Conversion in this population is likely magnified by competitive advantage for euploid cells during the transition to yeast-phase growth. Comparing the timing of morphological transitions of pure populations and the mixed population also shows that the presence of aneuploid cells affects the speed of the transition of euploid yeast to hyphae: at day 7, the mixed population is almost fully hyphal despite the presence of many euploid cells while at this timepoint the pure euploid cells are almost fully yeast.

### Chromosome 7 aneuploidy reduces virulence and proliferation in a murine model of infection

To uncover how Chr7 gain or loss affects virulence, we used the murine model of infection to compare the virulence of aneuploid strains, euploid strains, and a 1:1 mix of the two. We found that cells with an extra copy of Chr7 were drastically reduced in their ability to cause lethal infection compared to their euploid counterparts (Fig. 4A). Similarly, infections with aneuploid as opposed to euploid *Histoplasma* resulted in a significantly lower fungal burden by day seven both in the lung and spleen (Fig. 4B). When a 1:1 mix of aneuploid and euploid *Histoplasma* was used for infection, mouse survival was similar to that of mice infected with a pure euploid population (Fig. 4A). Interestingly, as the infection progressed, the percent of euploid cells significantly increased, a trend not found for yeast growing in vitro (Fig. 4C, S4A).

**Figure 4:**
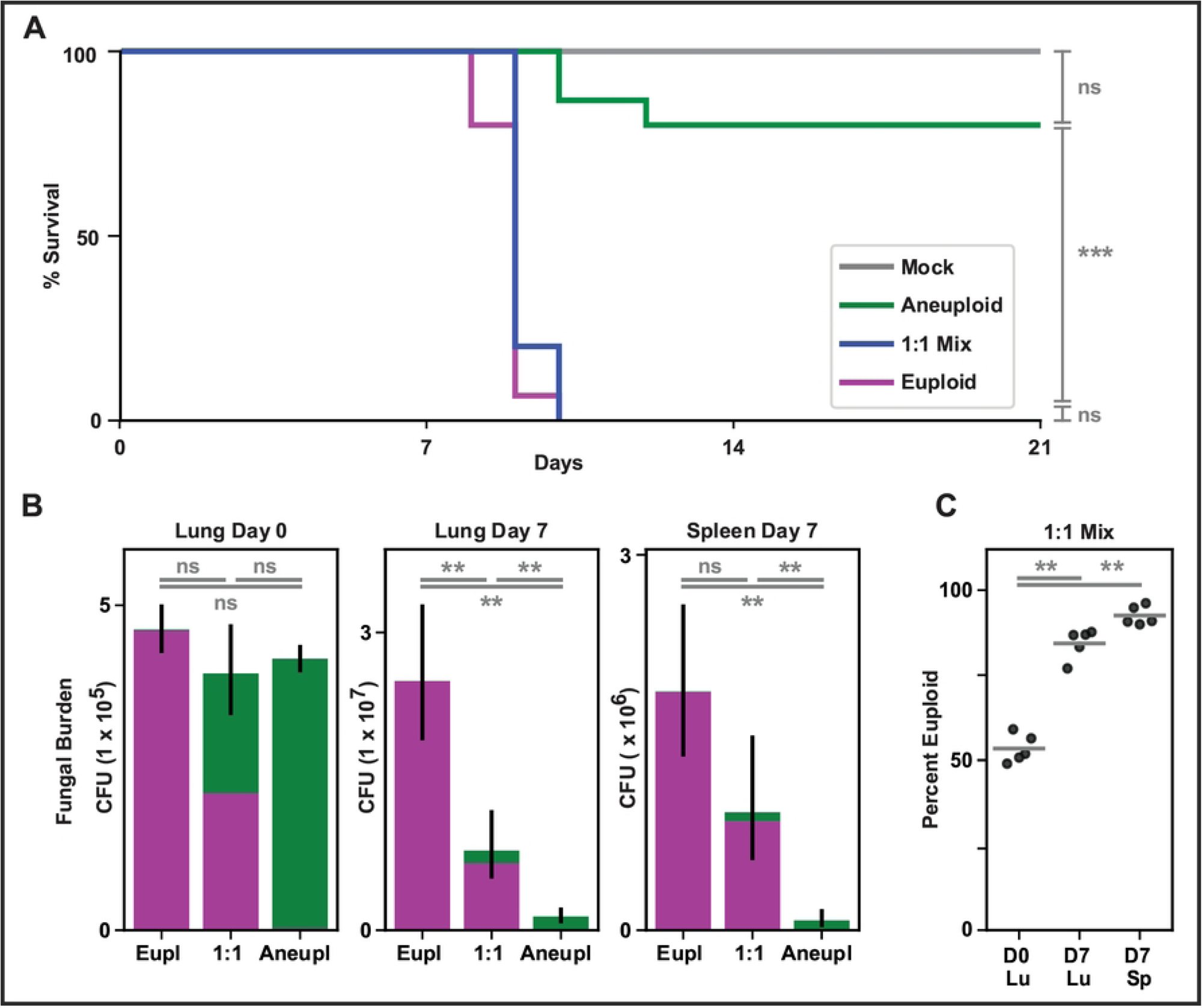
Aneuploidy reduces virulence in a mouse model. A) Survival curve of mice infected with aneuploid *Histoplasma* (green), euploid *Histoplasma* (magenta), a 1:1 mix of these two strains (blue), or mock infection (PBS, grey). Significance based on log rank test. B) Fungal burden by colony forming unit (CFU) in organs of mice infected with *Histoplasma*, with colony morphology assay used to determine ploidy of fungi obtained from organs. Portion of CFU encompassing each ploidy is indicated by color in bar graph, with aneuploid shown in green and euploid shown in magenta. Statistics on chart show significance by Wilcoxon test of total CFU between infection categories. C) Percent of fungal burden encompassing smooth (euploid) colonies for each replicate with mean shown by line. Statistics show significance by Wilcoxon test during 1:1 mix competition of portion of colonies that are euploid at Day 0 (hour 4 post-infection) in the lung versus portion that are euploid in lung (Lu) and in spleen (Sp) at Day 7.

### Yeast with a second copy of Chr7 display a hyphal-biased transcriptome

We next assessed the *Histoplasma* transcriptome for ploidy-dependent trends with an eye towards mechanisms by which the aneuploidy might affect morphology bias or virulence. We performed RNA sequencing of aneuploid or euploid yeast, aneuploid or euploid hyphae, and aneuploid or euploid cells at two and seven days into the transitions from yeast to hyphae or hyphae to yeast (Fig. S5).

By comparing steady-state hyphae and yeast, we observed that the global transcriptional signature of hyphae in comparison to yeast was consistent regardless of ploidy (Fig. 5A). Genes on Chr7 (shown in blue throughout Fig. 5) displayed similar distribution of expression in hyphae over yeast as the remainder of the genome (Fig. 5B). However, as expected, Chr7 transcripts were more abundant in aneuploid versus euploid cells (Fig. 5C-D), concordant with increased copy number of those genes. We compared the expression profile of aneuploid and euploid yeast, as well as aneuploid and euploid hyphae. Ploidy affected more transcripts in yeast (3178 transcripts with either increased or decreased abundance) than in hyphae (1513 transcripts).

**Figure 5:**
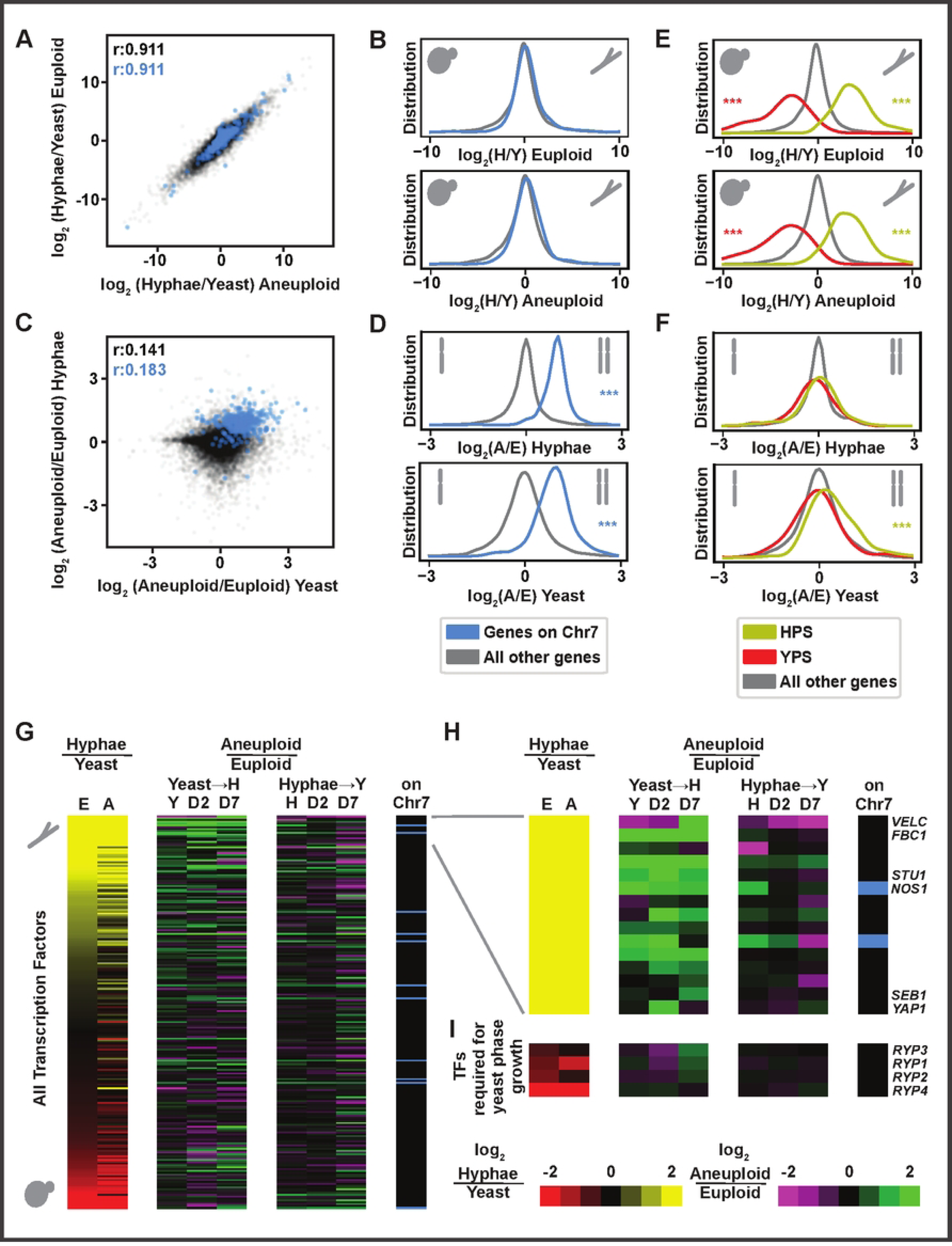
Aneuploid *Histoplasma* has a hyphal-biased transcriptome. A and C) scatter plots showing log_2_ ratios of transcript abundance. A) steady-state hyphae/steady-state yeast transcript ratios in aneuploid cells and in euploid cells. B) Aneuploid/Euploid transcript ratios in steady-state yeast and steady-state hyphae. Genes on chromosome 7 are shown in blue, all other transcripts are shown in grey. Pearson’s r correlation coefficients are shown in the upper left. B,D-H) Distribution of log_2_ ratios of transcript abundance, showing the same set of axes as used in A and B. Ratios from top to bottom include euploid hyphae/euploid yeast, aneuploid hyphae/aneuploid yeast, aneuploid hyphae/euploid hyphae, aneuploid yeast/euploid yeast. B,D) Transcripts encoded by genes on Chr7 are highlighted in blue. E-F) YPS transcripts are shown in red and HPS transcripts are shown in yellow. Asterisks are shown if significant by Wilcoxon test and if distribution median is least 5% from center to plot maximum or minimum. G) Heatmap showing all predicted transcription factors. The first two columns show transcript abundance in hyphae/yeast (column 1: euploid, column 2: aneuploid) (yellow versus red). Subsequent columns show transcript abundance in euploid versus aneuploid cells through yeast-to-hyphal transition (37°C to 25°C) and hyphal to yeast transition (25°C to 37°C) (green versus magenta). For each timepoint of each transition (steady state, day 2, and day 7), fit for each plotted transcript is derived from 3 replicates. Final column indicates transcripts encoded by genes on Chr7 in blue. H) Expanded view of the top 15 genes in G with named genes labeled. I) The same set of columns is shown for transcription factors required for yeast phase growth (*RYP1-4*).

We also analyzed differential expression for sets of previously defined *Histoplasma*-conserved yeast-phase specific transcripts (YPS) and hyphal-phase specific transcripts (HPS)^19^. We found that YPS had the expected increased abundance in yeast versus hyphae independent of ploidy just as HPS had the expected increased abundance in hyphae versus yeast (Fig. 5E). We found that HPS transcripts were also significantly more abundant in aneuploid versus euploid yeast (Fig. 5F). These data indicate that aneuploid yeast show a hyphal bias in their transcriptome, likely underlying the ability of these yeast to more rapidly undergo conversion to hyphae upon temperature shift.

We also assessed differential expression of TFs in this experiment based on a previously defined list of putative TFs^29^. We found that the TF transcripts that are most abundant in hyphae relative to yeast were often also more abundant in aneuploid yeast relative to euploid yeast, consistent with the hyphal bias of these cells (Fig. 5G-H). These TF transcripts showed even higher differential expression in aneuploid versus euploid cells during the transition from yeast to hyphae compared to steady-state yeast. Once aneuploid and euploid cells transition to steady-state hyphae, their transcriptomes look more similar and TF transcript abundance is no longer as differential. Similar trends were not observed in the hyphal to yeast transition (Fig. 5G, S5C), consistent with ploidy having a larger effect on yeast than on hyphae.

Two of the TFs that are upregulated in aneuploid cells, *STU1* and *FBC1*, neither of which are on Chr7, have been shown to be sufficient to induce the hyphal morphology when ectopically expressed in cells at 37°C^16^. However, we found that transcript abundance of the TFs required for yeast phase growth (*RYP1-4*^15,18^) were not strongly regulated by ploidy (Fig. 5I). Taken together, these data suggest that the transcriptome of aneuploid yeast renders these cells more primed for the hyphal transition compared to euploid yeast.

Although the YPS regulon is large, only a handful of genes have been tested for virulence in the mouse model. Five *Histoplasma* genes have been shown to affect mouse survival^30–33^, none of which are significantly regulated in aneuploid versus euploid yeast (Fig. S5D). However, among a recently defined set of 11 additional predicted virulence effectors^33^, 5 are significantly more expressed (mean 4.5 fold increase) in euploid versus aneuploid yeast (Fig. S5E).

### Identification of a transcription factor on chromosome 7 that promotes hyphal bias

Our analysis of matched yeast-biased and hyphal-biased strains allowed us to narrow down the region that was necessary for hyphal bias to a 334 kb critical region on Chr7 (Fig. 1C). This critical region contains 145 predicted genes (1.2 % of genes in the genome), including some with minimal evidence of expression and no predicted protein domains. None have been studied previously in *Histoplasma*. Through manual curation of the genes within this region (considering Pfam domain annotations, *Histoplasma* expression patterns, and annotated homologs), we generated five regions of interest, some containing >1 gene. The corresponding genes of interest included the only predicted TF in the region, two genes with distant homologues that affect morphology (protein kinase (PK) *YAK1* and chromatin modifying component *ADA2*^34–36)^, other PKs, and one gene encoding a small hyphal peptide (SHP) of unknown function with strongly hyphal-enriched transcription and translation^19^. Each of these five subregions was cloned into individual plasmids that were transformed into aneuploid and euploid *Histoplasma* strains (Fig. 6A, S6A-E) and the resultant effect on hyphal bias was determined.

**Figure 6:**
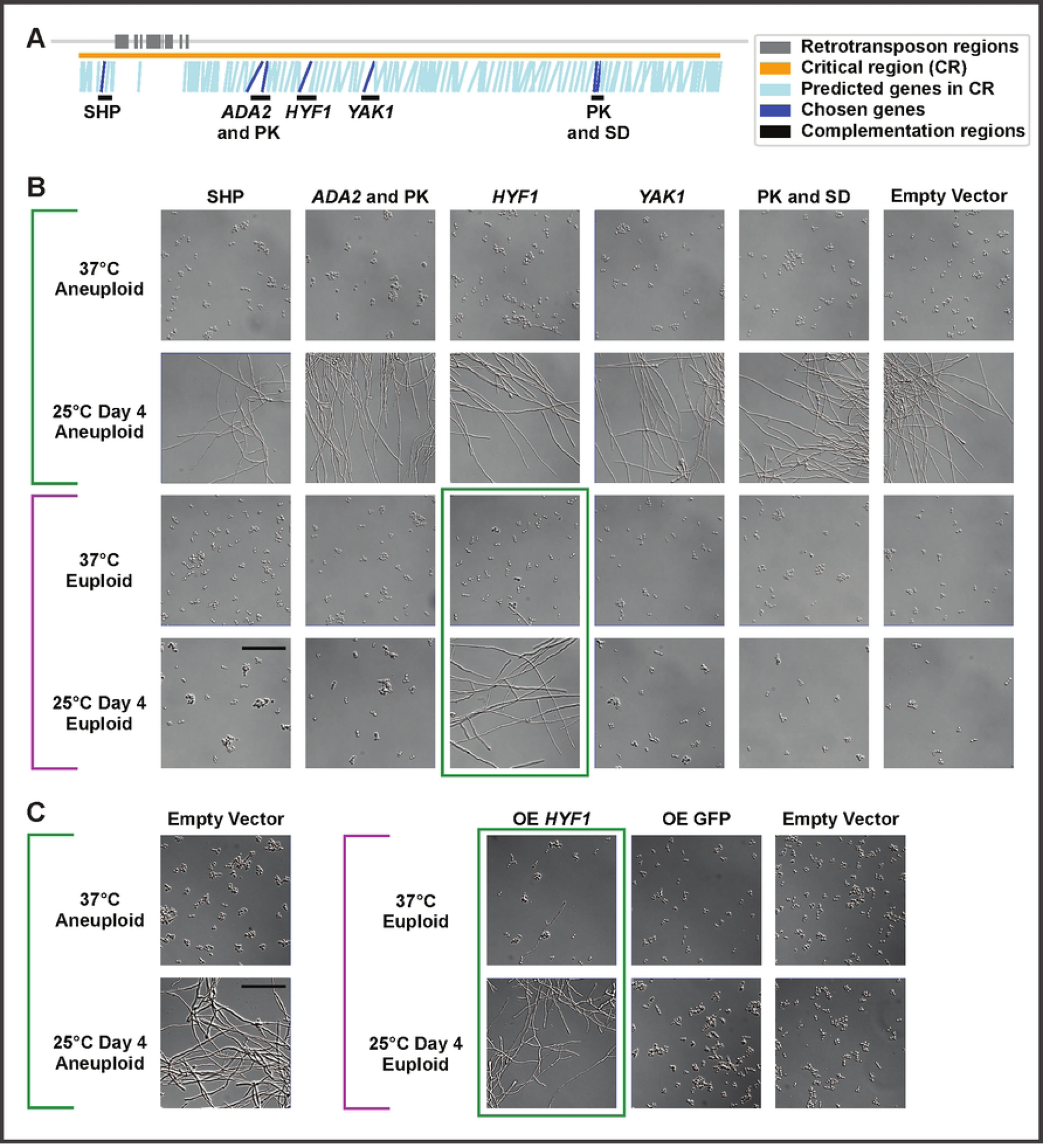
A chromosome 7 transcription factor, *HYF1*, is sufficient to confer hyphal bias. A) Locations of genes (blue) in the critical region (orange) with retrotransposon regions (grey) and regions chosen for increased copy number (black). Primary gene(s) chosen for each region are labeled and shown in dark blue. The critical region is 334 kb and contains 145 predicted genes. Chosen regions from left to right contain a small hyphal peptide (SHP), the chromatin modifying component *ADA2* (region also includes a PK), the TF *HYF1*, the PK *YAK1*, a region with a PK and a superoxide dismutase (SD). B) Microscopy of aneuploid and euploid cells at 37°C or after 4 days at 25°C with ectopic plasmids containing a portion of the critical region or an empty vector. Scale bar indicates 50 µm. C) Microscopy of aneuploid and euploid cells at 37°C or after 4 days at 25°C with overexpression (using the *ACT1* promoter) of *HYF1* or GFP (control) or an empty vector. Scale bar indicates 50 µm. Brackets and boxes in magenta and green indicate morphology bias.

Increasing the copy number of the critical region TF with its native regulatory elements was sufficient to confer hyphal bias in a euploid background (Fig. 6B). Average copy number of this plasmid in yeast was 1.4X based on normalized bulk sequencing coverage. The occasional hyphal cell was also observed at 37°C, concordant with this coverage average and the role of this TF in promoting hyphal growth. We named this TF *<u>Hy</u>phae Promoting <u>F</u>actor* (*HYF*) *1*. *HYF1* is orthologous to *CON7,* which affects morphology and virulence phenotypes in multiple fungal species^37–39^, and orthologous to *WOR4,* which drives the white-to-opaque transition in *Candida albicans*^40^ (Fig. S6F). Unfortunately, we were not able to successfully use CRISPRi to generate knockdown strains of *HYF1* despite several attempts. None of the other four gene segments conferred an increase in hyphal bias. We also overexpressed *HYF1* under the control of the *ACT1* promoter in euploid cells and observed strong hyphal bias, suggesting that this TF is a key regulator of hyphal development. (Fig. 6C).

### Increased copy number of *HYF1* replicates the transcriptome hyphal bias associated with increased copy number of Chr7

To determine if increased copy number of *HYF1* is sufficient to shift the transcriptome in the same manner as increased Chr7 copy number, we subjected euploid yeast carrying the *HYF1* plasmid to transcriptional profiling. Transcript abundance in *HYF1* plasmid versus empty vector control yeast was compared to transcript abundance in aneuploid versus euploid yeast. As expected, Chr7 CNV significantly affected the abundance of many more transcripts than increased *HYF1* copy number alone (Fig. 7A, B). Again as expected, a higher portion of transcripts with increased abundance in aneuploid cells versus transcripts up in *HYF1* are on Chr7 (30% versus 5%). 144 transcripts exhibited increased expression under conditions of both increased Chr7 copy number and increased *HYF1* expression. These transcripts were enriched for the previously defined HPS transcripts, which are highlighted in Fig. 7C. In contrast, the effect on the bulk of YPS transcripts was minimal (Fig. 7D).

**Figure 7:**
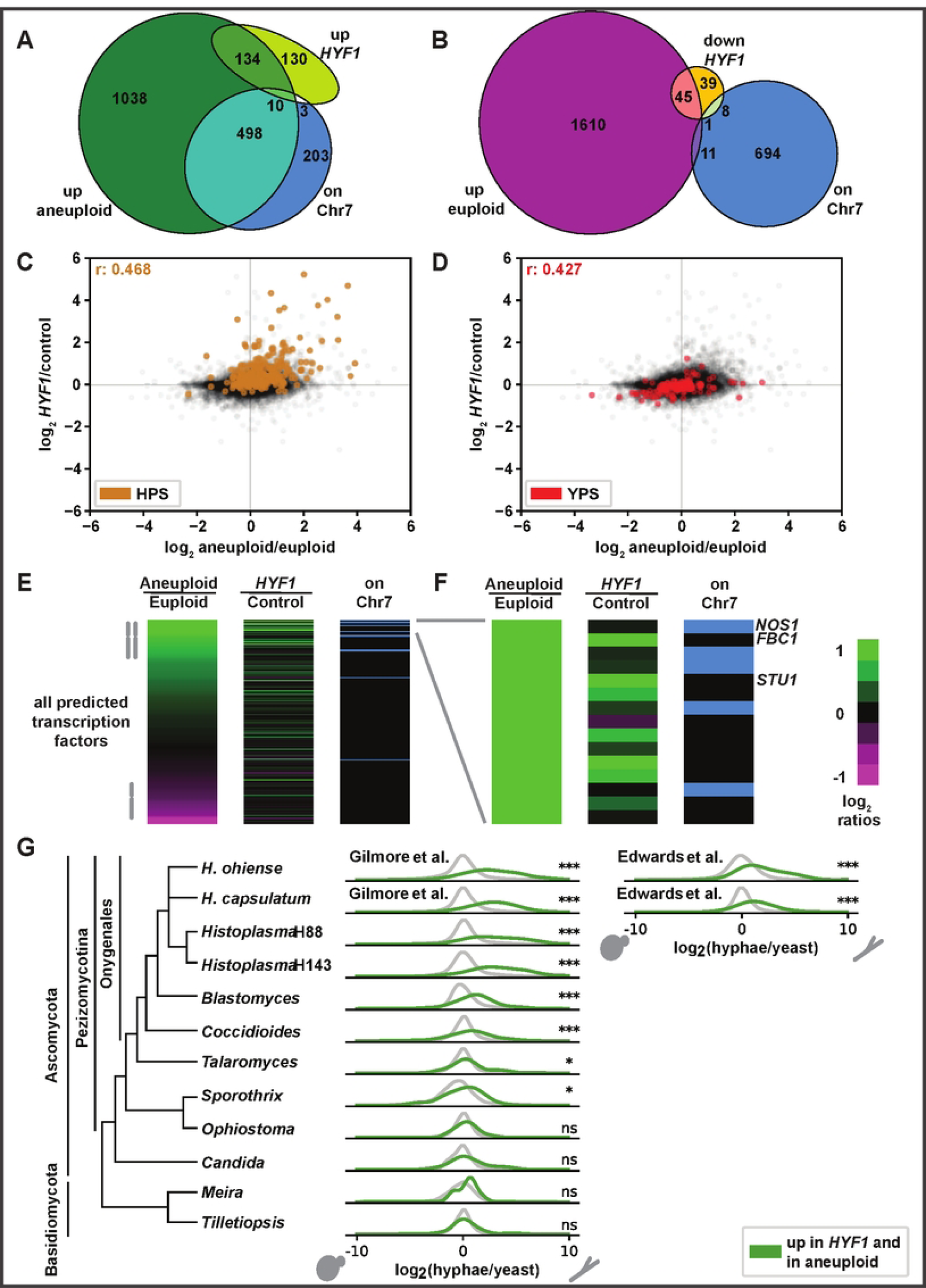
*HYF1* recapitulates some of the key transcriptional effects of aneuploidy. A) Venn diagram showing transcripts that reach significance and 1.5-fold increase in aneuploid versus euploid yeast (green) and *HYF1* increased copy number over control yeast (yellow-green) and genes on chromosome 7 (blue). B) Venn diagram showing transcripts that reach significance and 1.5-fold increase in euploid versus aneuploid yeast (magenta), 1.5-fold decrease in *HYF1* increased copy number versus control yeast (orange) and genes on chromosome 7 (blue). C-D) Scatter plots showing log_2_ ratios of transcripts in aneuploid versus euploid yeast (x-axis) and *HYF1* increased copy number over control yeast (y-axis) highlighting HPS genes (brown, in C) or YPS genes (red, in D). E) Heatmap showing all predicted TFs with the same two ratios as in C-D followed by a column indicating which genes are on Chr7 (blue). TFs are sorted based on the first column. F) The top 15 transcripts from E. G) Distribution of yeast versus hyphae transcript abundance in orthologs of all *Histoplasma* genes (grey) and genes up in *HYF1* and aneuploid (green). Asterisks indicate significance by Wilcoxon test.

Most TF transcripts with increased abundance in the aneuploid strain were either on Chr7 or instead had increased abundance in response to increased *HYF1* copy number (Fig. 7E). Of the 15 most abundant TFs in aneuploid vs euploid cells, 5 TFs were encoded on Chr7 and these TFs were not upregulated under conditions of *HYF1* increased copy number. In contrast, six of the remaining 10 TFs did show increased abundance under conditions of increased *HYF1* copy number (Fig. 7F). The two most differentially abundant TFs among these six, *FBC1* and *STU1*, have been previously shown to be sufficient to induce hyphal growth^16^. Increased expression of *STU1* and *FBC1* may be part of the mechanism by which increased copy number of *HYF1* induces hyphal bias.

It is possible that, while not required for hyphal bias, additional genes on Chr7 may also contribute hyphal phenotypes associated with aneuploidy. We also assessed the effects of increased copy number of an additional TF on this Chr7 found directly adjacent to the critical region that is orthologous to *RFEC (FFMA)* which is required for normal hyphal growth in *Aspergillus*^41^. This TF, which we name *HYF2,* slightly increases some of the morphological and transcriptomic phenotypes of cells carrying a second copy of Chr7 (Fig. S7). However, its effect on both morphology and transcriptome is much more subtle than that of *HYF1*.

These data defined a *HYF1* regulon of transcripts whose abundance changed in response to increase in *HYF1* copy number.^19,29,42–50^ To determine if the *HYF1* regulon is broadly correlated with filamentous growth in other fungi, we assessed the morphology-specific expression of these genes in available fungal yeast versus hyphae datasets (Fig. 7G)^19,29,44–50^. Within the thermally dimorphic pathogens in pezizomycotina, there is significant enrichment of *HYF1* regulon orthologs among genes up in hyphae. In contrast, we do not observe this enrichment for the close *Sporothrix* relative *Ophiostoma novo ulmi*, a plant pathogen with temperature-independent dimorphism. Likewise, the enrichment does not hold for the more distantly related ascomycete *Candida albicans*, a mammalian pathogen with a 37°C induced yeast to hyphal transition, nor for the basidiomycete plant pathogens *Meira miltonrushii* and *Tilletiopsis washingtonensis* which, like *Ophiostoma*, have temperature-independent dimorphism.

Notable genes with conserved hyphal expression in most or all of the thermally dimorphic fungi include the ortholog of A. nidulans developmental regulator *esdC*^51^, the ortholog of *Metarhizium anisopliae* cold shock protein *CRP2*^52^, and the transcription factors *FBC1* and *STU1*. The conserved hyphal expression for *STU1* in particular is striking. In is up at least 2x in hyphae for all of the pezizomycotina, with only *Ophiostoma* failing to pass the 5% FDR significance criterion, and statistically significant but, at 1.99 X, just under the fold change criterion in Meira.

As most of the species with conserved hyphal expression of the *HYF1* regulon are both thermally dimorphic pathogens of mammals and also the more closely related to *Histoplasma*, we can not distinguish whether this expression pattern is a feature of this class of pathogens versus a conserved property of pezizomycotina versus a more general pattern that is obscured in more distant relatives of *Histoplasma* due to increased difficulty of ortholog mapping. Nevertheless, the stronger hyphal enrichment of the *HYF1* regulon in *Sporothrix* relative to *Ophiostoma* suggests that this enrichment may be a property of thermal dimorphs.

## Discussion

We discover here that gain or loss of an extra copy of a particular *Histoplasma* chromosome, a reversible genetic change, has profound effects on the switch between the environmental and host forms as well as the ability of the organism to thrive during infection. In the laboratory, gain or loss of this chromosome occurs at the rate of 5-11×10^-5^ per generation, and our detection of CNV of this chromosome in 16% of previously sequenced natural isolates indicates that it occurs broadly. Notably, euploid cells have a significant competitive advantage in the host whereas cells with an extra copy of this chromosome have an enhanced ability to switch to the environmental form in response to temperature cues.

Facile gain and loss of a specific *Histoplasma* chromosome may benefit *Histoplasma* by increasing phenotypic variability, helping populations survive abrupt transitions between environment and host (Fig. 8). It may be beneficial for a subpopulation of cells to be primed to switch morphology while another subpopulation is recalcitrant to fluctuations in stimuli including temperature, which can overlap between human airways and the soil^53^. Hyphal cells are occasionally observed in human tissues in cases of severe histoplasmosis, and it is possible that this morphological diversity could be evolutionarily beneficial to *Histoplasma* in preparation for possible transition to environmental growth^54–56^. We hypothesize that Chr7 aneuploidy may act as a bet-hedging system, allowing variability in the morphological response. We also find it interesting that in mixed aneuploid-euploid cultures transitioning from yeast to hyphae, the presence of aneuploid cells seems to promote euploid cells to transition from yeast to hyphae more rapidly. It is possible that in addition to functioning as a bet-hedging system, successful hyphal growth of a subpopulation may act as a bellwether, inducing morphological transition in more of the population, *e.g.,* by production of a secreted factor that promotes hyphal growth.

**Figure 8:**
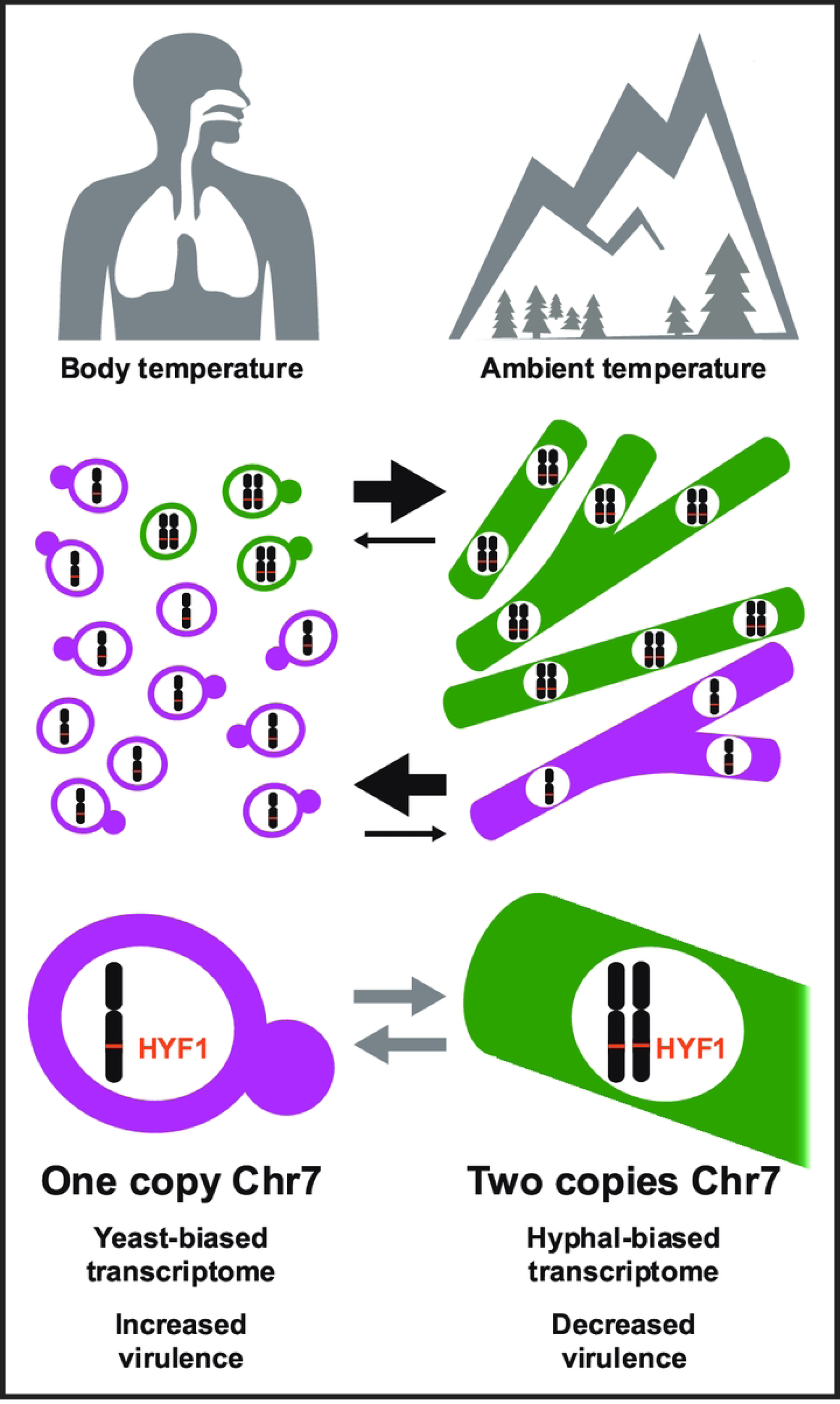
Rapid gain and loss of Chr7 aneuploidy may increase phenotypic diversity. Our data suggest that rapid gain and loss of the Chr7 aneuploidy may benefit *Histoplasma* by rapidly increasing phenotypic diversity, helping populations survive frequent and abrupt transitions between environment and host. *Histoplasma* grows as yeast in the mammalian body and in the laboratory when grown at 37°C but as hyphae in the environment or in the laboratory when grown at 25°C. Cells with a second copy of chromosome 7 are biased towards hyphal growth and outcompete euploid cells in the yeast-to-hyphal transition (black arrows). Euploid cells (with one copy of each chromosome) are biased towards yeast growth and outcompete in the hyphal to yeast transition (black arrows). Cells frequently gain and lose a second copy of Chr7 (grey arrows). Cells with one copy of Chr7 have increased virulence in comparison to cells with two copies of the chromosome. Cells with two copies of Chr7 have a hyphal-biased transcriptome as do cells with increased copy number of *HYF1*, a TF on Chr7.

Since this CNV is common in sequenced natural isolates and prevalent in lab strains, it is intriguing to speculate that the aneuploidy could have influenced prior experimental findings relevant to *Histoplasma* virulence or filamentation. At the minimum, this CNV explains the phenotypes previously attributed to *MSB2* in *Histoplasma*^20,21^. While ploidy variation is clearly observed in human clinical isolates of *Histoplasma*, we do not yet know if *Histoplasma* ploidy variation correlates with clinical presentation or is influenced by the immune status of the host. Exceedingly few environmental isolates are available, so it remains to be seen the level at which ploidy variation is present in *Histoplasma* growing in the environment.

Our work revealed that aneuploidies in *Histoplasma* are by far most common on Chr7 versus other chromosomes. In contrast to *Histoplasma*, the incidence of aneuploidy is often spread more evenly between chromosomes in model species although certain aneuploidies are associated with adaptation to specific stressors^26,27,57–59^. While it is possible that relative rate of Chr7 aneuploid acquisition (e.g. increased nondisjunction of Chr7 versus other chromosomes) may contribute to the frequency at which we observe Chr7 CNV, we hypothesize that the primary driver is relative aneuploid fitness burden among the seven chromosomes. For both *H. ohiense* and *H. mississippiense*, the aneuploid chromosome contains the fewest genes, which could be related to a lower fitness cost^60^. However, partial chromosome CNV is also most common within Chr7. We thus hypothesize that minimal fitness cost of duplication of genes on Chr7 and conditional competitive advantage of this CNV are the primary drivers of its frequency.

Identification of the critical region facilitated our discovery of the TFs *HYF1* and *HYF2* as hyphae-promoting factors. While these genes have not been investigated previously in *Histoplasma*, each is orthologous to a gene with functions related to morphology and cell fate. *HYF1* is orthologous to the *C. albicans* TF *WOR4*, which is involved in a transcriptional network regulating white-opaque switching^40^. *Histoplasma* orthologs of other genes within this *Candida* network are also involved in the *Histoplasma* regulation of morphology, including *RYP1*, an ortholog of *Candida WOR1*, and *STU1*, an ortholog of *Candida EFG1*^15,40^. *HYF2* is orthologous to the *Aspergillus fumigatus* protein *FFMA*, whose deletion results in reduced growth^41^. Thus far we have not been able to successfully generate knockdown strains targeting *HYF1* or *HYF2*. Genetic manipulation can be challenging in *Histoplasma*, so it is unclear if the lack of a knockdown strain indicates that, like *FFMA, HYF1* and *HYF2* are required for normal growth.

The effect of Chr7 copy number variation was more evident in yeast than in hyphae. Aneuploid yeast transitioned to hyphae much more quickly than euploid yeast transitioned to hyphae. However, aneuploid and euploid hyphae demonstrated only minor differences in speed of conversion to yeast. Similarly, while we observed a hyphal bias in the transcriptome of aneuploid yeast, suggesting that they were primed to transition more quickly to hyphae, euploid hyphae did not display as much yeast bias in their transcriptome. It is possible that some of the observed relative effect of ploidy may be attributable to our better understanding of the yeast-to-hyphal transition than our understanding of the hyphal-to-yeast transition. The yeast-to-hyphal transition requires less time than the reverse transition and is more synchronous under current culture conditions, and is thus more commonly performed in the laboratory. In fact, this is the first publication of transcriptional profiling of hyphal cells shifted to yeast-inducing conditions, providing a dataset that will be relevant to subsequent investigations of thermal dimorphism.

Our observation that a reversible chromosomal aneuploidy has profound effects on morphologic transitions and disease is essential to understanding thermal dimorphism and pathogenesis in this important group of understudied human pathogens. Although epigenetic changes that drive switches are more commonly reported, specific reversible genetic changes akin to this *Histoplasma* aneuploidy have been observed previously^61–64^. Such changes include inversion of DNA sequences, homologous recombination between transposons, and aneuploidies, and likely often have an intermediate stability between that of epigenetic modifications and standard genetic mutations. These reversible genetic changes often confer benefits in a subset of the ecological or pathogenic niches inhabited by a microbe. We posit that similar yet unidentified genetic changes are likely important to the pathogenesis of many additional microbes, conferring semi-transient diversity to improve population survival through repeatedly fluctuating conditions.

---

**Supplemental Table S1** Strains used

**Supplemental Table S3.**
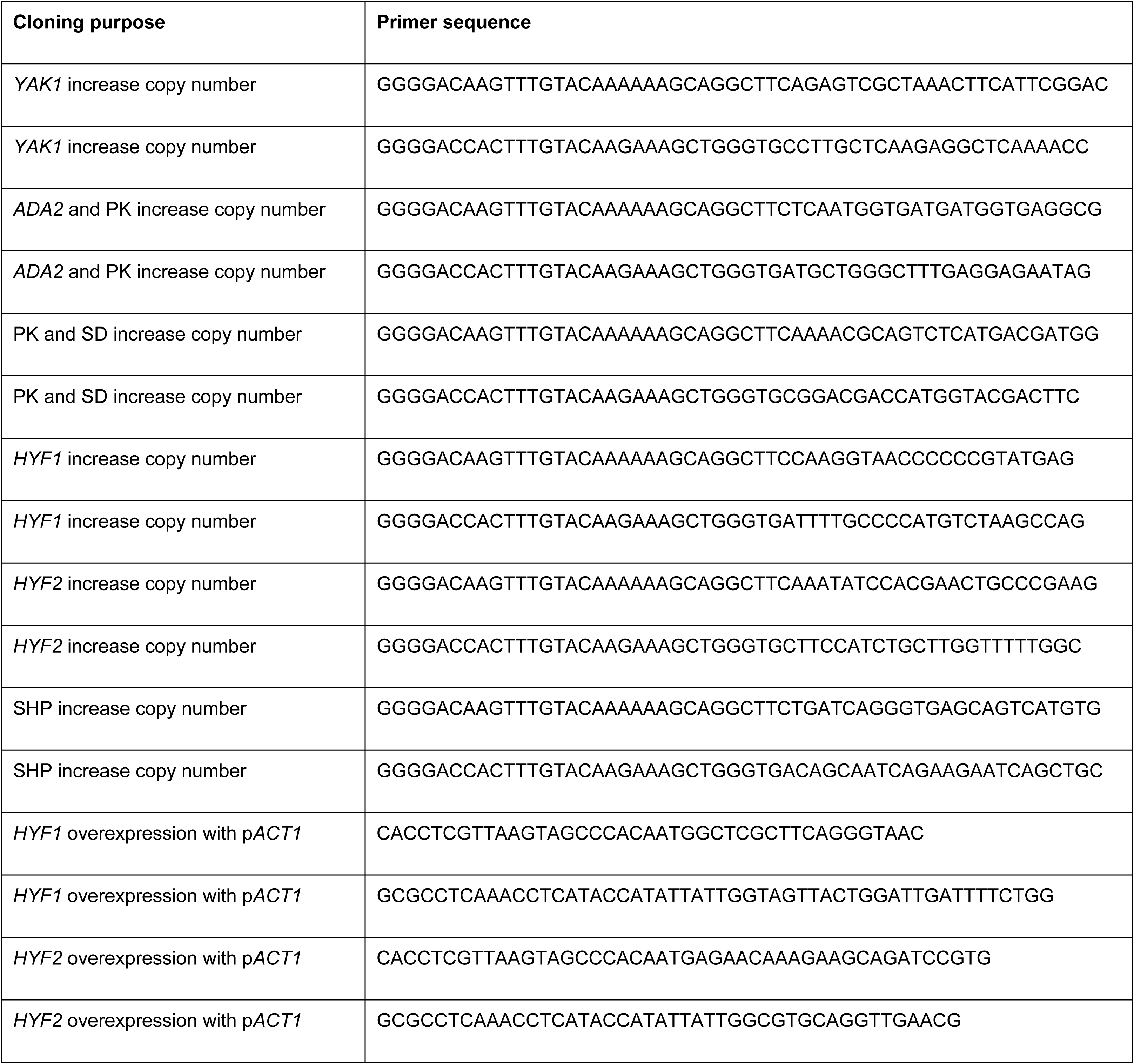
Primers used for cloning.

**Supplemental Table S3:** KALLISTO estimated counts and LIMMA-fit values for time course expression profiles.

Excel-compatible tab-delimited text conforming to JavaTreeView extended CDT format. Each row is a transcript, with the UNIQID column giving the ucsf_hc.01_1.G217B systematic gene name.

The NAME column gives short names taken from Data S1 of Voorhies, 2022^65^ with additional names and corrections based on human curation. The next 36 columns give KALLISTO estimated counts for each transcript in each sample. The next columns give LIMMA BH-adjusted p-values for differential expression in each of the 20 contrasts, and the corresponding LIMMA-fit log2 ratios are given in the final 20 columns. Description and HcG217B_pred (WUSTL G217B predicted gene accessions), HcG217B_rc (Edwards et al G217B transcript accessions^44^), and HcG217B_acc (UCSF1 gene accessions^65^); and Ryp1_ChIP, Ryp2_ChIP, Ryp3_ChIP, and Ryp4_C (indicating genes with promoters bound by Ryp TFs in Beyhan, 2013^17^) are taken from Data S1 of Voorhies, 2022^65^. UCSF2_acc and UCSF3_acc give gene accessions from the G217B assemblies of Heater, 2025^22^. CR is “critical_region” for genes in the critical region, “chr7” for genes on chromosome 7 but outside of the critical region, and blank for all other genes. GWEIGHT is a place-holder column for JavaTreeView compatibility. The estimated counts in this file are sufficient to recapitulate the LIMMA analysis.

**Supplemental Table S4**: KALLISTO estimated counts and LIMMA-fit values for expression profiles of strains with elevated HYF1 or HYF2.

Excel-compatible tab-delimited text conforming to JavaTreeView extended CDT format. Columns are as in table for time course expression profiles, except that the count, p-value, and log2 ratio columns correspond to the HYF1 and HYF2 elevated expression experiment. The estimated counts in this file are sufficient to recapitulate the LIMMA analysis.

**Supplemental Table S5**: Data sources for comparative transcriptome analysis.

Excel-compatible tab-delimited text. Each row corresponds to a profile plotted in Fig 7G. Columns give a shorthand name, taxonomic details (genus, species, strain), data accessions (GEO and SRA) and reference (PubMed PMID), sequencing layout (paired or single), extra flags supplied to KALLISTO (see Materials and methods), the specific SRA sample accessions used for the yeast or hyphae expression profiles, and the source (URL or GenBank accession) of the transcriptome sequences used to build the KALLISTO index.

**Supplemental Table S6**: Comparative transcriptome profiles.

Excel-compatible tab-delimited text conforming to JavaTreeView extended CDT format. UNIQID, NAME, Description, and annotation columns are as for supplemental table S3. Remaining columns, distinguished by the shorthand names from table S5, give ortholog gene IDs, LIMMA estimated log ratios, and LIMMA p-values for each comparison dataset.

**Supplemental Table S7**: Comparative transcriptome profiles for genes up in *HYF1* and aneuploid.

Excel-compatible tab-delimited text conforming to JavaTreeView extended CDT format. Identical to table S6, but restricted to genes up in *HYF1* and aneuploid (green curves in figure 7G).

## Supplemental figure legends

**Supplemental Figure S1:**
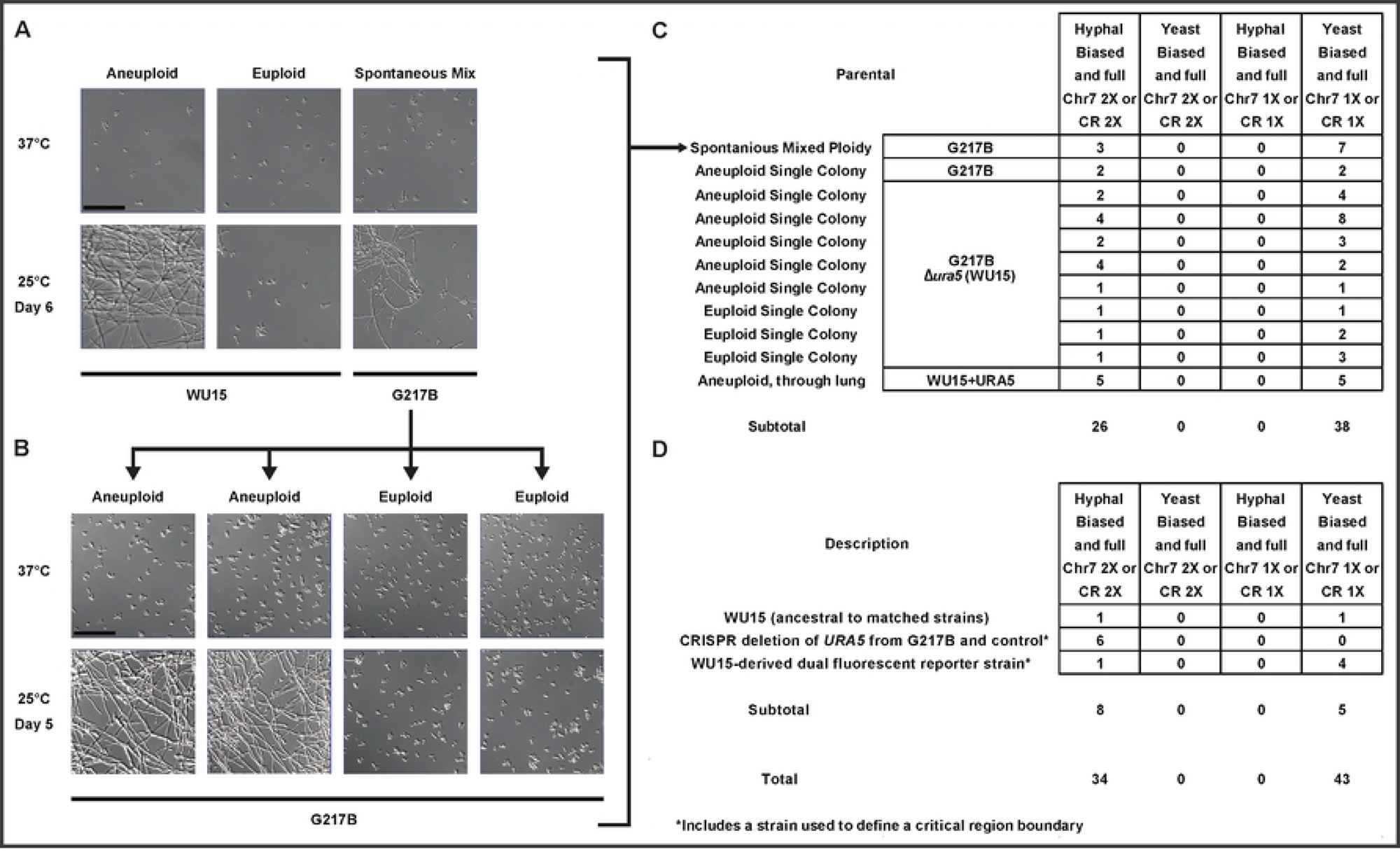
A) Microscopy of strains at 37°C and after 6 days at 25°C. A population of wildtype *Histoplasma* spontaneously obtained an intermediate morphological phenotype. Ten random single colonies were isolated from this population, three of which were found to have a hyphal-bias and seven of which were found to have a yeast-bias. B) Four representative isolates are shown. These ten isolates were subsequently sequenced, and a perfect correlation between ploidy and morphology bias was found, as shown in the first row of data in C. C) Summary of the sources for 64 strains for which paired aneuploid and euploid strains were derived from a parental source (64 strains shown in Fig. 1D). For these strains, at least one matched strain was generated at the same time, from the same parental source, but of the opposite morphology bias. The first row is described in S1A-B. For subsequent rows, colony morphology-bias was used to isolate yeast-biased and hyphal-biased progeny of aneuploid cells, euploid cells, and cells from a mouse lung. D) Sources for the 13 strains that were included in the analysis in Fig. 1C. These include strains ancestral to some of those shown above, as well as strain sets used to define a boundary of the critical region (indicated by asterisks) and controls. A total of 77 strains were phenotyped then subsequently genotyped for this analysis. Fisher exact p-value was < 1×10^-7^ for correspondence between morphology bias and critical region coverage. More information on strains is available in supplemental table 1.

**Supplemental Figure S2**: Heatmaps showing copy number (CN) variation in natural isolates (clinical isolates and a limited number of available environmental population isolates) of *H. ohiense* (A) *H. mississippiense* (B) and other *Histoplasma* (C). The number of isolates in each category is listed parenthetically on the y-axis. The *H. ohiense* reference genome is used for other *Histoplasma* isolates shown in C. For each CNV heatmap, the critical region for each genome is highlighted in orange. Chromosomes are indicated, along with location of rDNA (R).

**Supplemental Figure S3**: A) Optical density (OD_600_) growth curve of aneuploid and euploid yeast grown in standard yeast liquid growth conditions over the course of 141.5 hours. Fill indicates standard deviation at each point. B) Plots showing copy number variation in strains selected for sequencing from aneuploid gain and loss rate experiments, which are also included in the counts in Figure 1C-D and S1.

**Supplemental Figure S4**: A) Competition of aneuploid and euploid yeast grown continuously for four weeks showing fraction of aneuploid starting strain barcode from full genome sequencing. Fill indicates standard deviation at each point.

**Supplemental Figure S5**: A-B) Distribution of log ratios for expression in aneuploid versus euploid cells as in Fig. 5 showing (from top to bottom) steady state yeast, a 2-day transition to hyphae, a 7-day transition to hyphae, steady state hyphae, a 2-day transition to yeast, and a 7-day transition to yeast. Distributions of genes in chromosome 7 are shown in blue in A, HPS and YPS gene distributions are shown in B. C) Heatmap with the same columns as in Fig. 5 I-L showing all predicted genes. Genes are sorted based on the ratio of the first and third data columns, as noted by arrow heads below heatmap. (Specifically, the sort is on the element-wise arctangent of (euploid hyphae/yeast)/(yeast aneuploid/euploid) as implemented in arctan2 from numpy). Approximate locations of genes of interest are noted to the right of this heatmap. D) Heatmap with the same columns as showing genes that have been shown to affect mouse survival. E) Heatmap with the showing previously defined set of additional virulence factors.

**Supplemental Figure S6**: A-E) Locations chosen for regions of the critical region added on ectopic plasmids (black bars). Each plot in A-F shows ribosomal footprinting^19^ in yeast (red) and hyphae (green) above RNA-seq in yeast (red) and hyphae (olive) followed by locations of predicted genes (grey) then *RYP1-4* ChIP enrichment in (red, green, blue, and purple, respectively)^17^. For each plot, primary gene is labeled above plot and shown in darker grey fill while adjacent upstream gene is labeled below plot and indicated by black line in predicted gene subplot. D) The region used for overexpression of *HYF1* (on the ACT1 promoter) is also indicated by grey bar. Y-axis limits are consistent in A-E. F) Multiple alignment of *HYF1* and orthologs, colored by conservation and annotated as in Fig S1 of Odenbach et al^66^.

**Supplemental Figure S7**: A) Location of *HYF2* and complementation region as shown for other genes in Fig. 6A. B) Microscopy of cells with increased copy number of *HYF2* as shown for other genes in Fig. 6B. C) Microscopy of cells with overexpression of *HYF2* driven by the *ACT1* promoter as shown for *HYF1* in Fig. 6C. Experiments were performed simultaneously with those shown in Fig. 6 and thus share control images. D) Locations of regions used for increased copy number of *HYF2* and overexpression of *HYF2* on the *ACT1* promoter as shown for other genes in Fig. S6A-E. Y-axis limits are the same as in Fig. S6A-E except the ChIP track which has a maximum y value of twice that shown in Fig. S6 to show full range. E) Heatmap showing all predicted transcripts with the same columns as in Figure 5 in addition to a column showing transcript abundance in *HYF2* increased copy number yeast over yeast containing a control plasmid. Transcripts are sorted based on the first column (aneuploid/euploid). F-K) Plots showing transcriptomic data for *HYF2* as shown for *HYF1* in Fig. 7A-F. Note for H-I the y-axis change in scale used to show the more subtle effects of *HYF2*. L) Multiple alignment of *HYF2* and orthologs, colored by conservation.

## Methods

### Ethics statement

All animal work was approved under UCSF Institutional Animal Care and Use Committee protocol AN197403.

### *Histoplasma* growth conditions

Yeast were grown at 37°C with 5% CO_2_, and hyphae were grown at 25°C without CO_2_. Culture of hyphae was performed in a Biosafety Level 3 facility. For liquid growth, cells were grown in *Histoplasma*-macrophage medium (HMM) in an orbital shaker at 120-150 RPM. During liquid growth, cells were passaged at a 1:25 dilution every 2-3 days for yeast and weekly for hyphae. For growth on plates, HMM agarose was used. Plates and liquid HMM were supplemented with 0.2 mg/mL uracil for growth of uracil auxotroph strains. The glucose in HMM media was replaced with equimolar N-acetyl-glucosamine (GlcNAc) in indicated experiments to increase the speed of transition from yeast to hyphae^67^.

### Histoplasma strains

Strains used to assess morphology-CNV correlations are derived from clinical isolates G217B, CI_4, CI_9, CI_43, and UCLA as described for Fig. 1 and Fig. 2. The *URA5* deletion strain WU15^68^, which was derived from *H. ohiense* clinical isolate G217B is otherwise used unless noted. This includes otherwise isogenic pairs of aneuploid and euploid strains used for experiments shown in Fig. 1A-B, Fig. 2E-H, Fig. 3, and Fig. 5. WU15+*URA5* is used for mouse infections shown in Fig. 4. One non-genic SNP was present between euploid and aneuploid strains used in this experiment. Otherwise isogenic pairs of aneuploid and euploid WU-15 strains were transformed with plasmids to obtain strains used in Fig. 6 and Fig. 7 as described below. Strains used in this paper are also described in supplemental table 1.

### Assessment of morphology and morphology bias categorization from colony appearance and microscopy

To generate single colonies to assess colony appearance, cells in a yeast culture were briefly sonicated, counted with a hemocytometer, then plated to obtain around 40-100 colonies per plate. Plates were grown at 37°C and 5% CO_2_ until colonies were small but clearly visible, around one week, then moved to 25°C without CO_2_. Colonies were allowed to grow until the morphology differences between yeast-biased and hyphal-biased controls were stark, around ten additional days. For categorization of morphology bias by microscopy of liquid cultures, yeast cultures were passaged 1:25 into HMM+GlcNAc on day -2 and then on day 0 were passaged to OD_600_ 0.2 in HMM+GlcNAc and then moved from 37°C to 25°C. Microscopy samples were taken at subsequent days as indicated for individual experiments and images. Samples for morphology bias categorization were taken between days 2 and 10 when sharply delineated morphology differences were consistently observable for control strains. Yeast-biased isolates had morphology scores of 1 (all yeast) or 2 (vast majority yeast) while hyphal-biased isolates had morphology scores of 4 (vast majority hyphae) or 5 (all hyphae). For imaging, cells were fixed in 4% PFA for 30 minutes and stored at 4°C until DIC imaging on a Zeiss AxioCam MRM microscope.

### DNA sequencing and analysis, including ploidy categorization

Genomic DNA extractions and sequencing were performed as previously described^22^. All DNA extractions other than for the five dual fluorescent strains were performed using the bead beating protocol^22^. Genomic DNA extraction for the five dual fluorescent strains were performed using the slightly modified protocol based on the Qiagen Gentra Puregene Yeast/Bacteria kit (158567)^22^. Illumina sequencing was performed at UCSF Center for Advanced Technology, the Chan Zuckerberg Biohub – San Francisco, or SeqCenter, LLC (Pittsburgh, PA) . Reads were aligned to assembly UCSF3 for non *H. mississippiense* strains and to the WU24 reference assembly for *H. mississippiense* strains.

Coverage per base was normalized based on median coverage per base between the genes SRE1 and MET1, Chr3 bases 2533970 to 3746857 for UCSF3 and the syntenic region on Chr2 for assembly WU24, 2400021 to 2941628. Strains sequenced from in-lab experimentation were generated directly from single colonies and had distinct nearly 1X or 2X coverage throughout nonrepetitive chromosomal regions. For these isolates, coverage through the critical region determined ploidy categorization. One boundary of the critical region was defined by the CNV region in two strains derived by CRISPR deletion of URA5 from G217B. These two strains retain hyphal bias despite having a very limited region of Chr7 duplication. The other boundary was determined by the CNV in four yeast biased strains that have duplication of only the left-hand portion of Chr7. These four strains were selected for yeast bias versus their ancestral aneuploid strain, a hyphal biased strain containing fluorescent reporters. The critical region was found to be Chr7 bases 1466188 to 1800070 in assembly UCSF3 and Chr6 bases 234820 to 597144 in assembly WU24. For population isolates, which included a few with intermediate coverage levels, 1.25X median coverage in the critical region was used as a threshold to determine ploidy categorization.

### Assessment of the rate of aneuploidy gain and loss

For determination of colony morphology bias conversion rate, proliferation from a single parental cell was counted by hemocytometer through growth in colonies and brief growth in liquid culture for an average of 31 doublings total prior to the assessment of progeny morphology. To determine aneuploid loss rates, a total of 11364 colonies were categorized for morphology with 44 colonies switching in morphology among 6 experimental replicates (replicates averaged 32 doublings). To determine aneuploid gain rates, a total of 9750 colonies were categorized for morphology with 19 colonies switching in morphology among 5 experimental replicates (replicates averaged 31 doublings). In these rate calculations, the few sectored colonies (*e.g.*, colony adjacent to green star in Fig. 1B) were categorized by the dominant morphology of the colony.

### Conditions for competition assays

To obtain matched steady-state yeast and hyphae, cultures were started from single yeast colonies and grown as yeast for two days. These starter cultures were used to inoculate plates without GlcNAc at 37°C and with GlcNAc at 25°C which were grown in parallel for 20 days. Hyphal cultures were then started from 25°C plates, grown for 2 days in liquid HMM supplemented with GlcNAc to ensure full hyphal conversion, passaged 1:6 in HMM without GlcNAc for five days, full hyphal growth was confirmed by microscopy, and hyphae were passaged 1:25 in media without GlcNAc and grown for seven days. Yeast cultures were started from 37°C plates and passaged 1:25 3 times a week in media without GlcNAc for the same two weeks as the hyphal cultures. After 2 weeks of steady state liquid growth, competition mixtures were made. Yeast were mixed at equal OD_600_ (where appropriate) and diluted to OD_600_ of 0.2, and hyphae were mixed to equal volume (where appropriate) and total dilution of 1:25. For steady-state yeast competition, cells stayed in standard yeast conditions with passage three times a week. For morphology transitions, yeast were then brought to standard hyphal conditions and hyphae were brought to standard yeast conditions on day 0, and cultures were passaged 1:25 every seven days throughout the competition, at which points samples for DNA sequencing and microscopy were taken as described above.

For competition experiments, at least four microscopy images were taken per flask per timepoint. All images were combined and randomized (with 25 repeating images to ensure consistency) then categorized by morphology for morphology score as previously described^22^. Images were randomly shown and categorized based on an image key as all yeast (1), vast majority yeast (2), mixed (3), vast majority hyphae (4), all hyphae (5).

In competition assays, two barcodes, defined as the alleles at two non-coding SNPs, were used to identify the starting strains as either euploid or aneuploid. Each barcode was assessed for either euploid or aneuploid starting cells to ensure that the SNP barcode itself did not affect the data. Fraction aneuploid barcode was calculated at each SNP as (aneuploid barcode allele counts)/(total counts for both alleles) and the mean over the two locations is reported. Chr7 copy number was calculated based on median coverage in Chr7 versus median coverage in the normalizer region.

### Assessment of survival and fungal burden in murine model

Murine infections were performed as previously described^33^ using 8-12 week-old female C57Bl/6 (Jackson Laboratories Strain 000664) mice infected intranasally with 1.0 x 10^6^ WU15+*URA5* yeast per mouse. For survival analysis, mice were monitored daily for both symptoms of disease (hunching, panting, ears tucked back) and weight loss (weight loss relative to day 0) as part of our euthanasia criteria over the course of 21 days. Mice were euthanized after they exhibited 3 days of sustained weight loss to ≤ 75% of their maximum weight in addition to one other symptom. To determine fungal burden and to count and categorize colonies of each morphology bias, lungs and spleens were collected from infected mice and homogenized in PBS. Homogenates were sonicated, diluted, plated on HMM and grown at 37°C then at 25°C for 8-13 days, at which point colonies with each phenotype were counted as previously described.

### RNA sequencing

Steady-state and transitioned cultures were prepared as for competition assays. Steady state and two-day transition timepoints were taken from two-day old cultures. Seven-day transition cultures were collected after seven days in conditions inducing the opposite morphology without passaging. Each condition was assessed in biological triplicate; media was not supplemented with GlcNAc for any of these conditions. *HYF1* increased copy number, *HYF2* increased copy number, and control samples for RNA sequencing were collected from cultures started at OD_600_ of 0.2 then grown for 2 days in standard yeast conditions in HMM media not supplemented with GlcNAc, the same as was done for aneuploid and euploid steady state yeast included in the RNA sequencing transition assays. Cells were collected by filtration, flash frozen in liquid nitrogen, and stored at -80°C prior to RNA extraction. RNA extraction, mRNA isolation, and RNAseq library preparation were performed as previously described^16^. Average fragment size and presence of excess adapter was determined with High Sensitivity DNA Bioanalyzer chip from Agilent Technologies (Santa Clara, CA). Library concentration was measured by Qubit™ dsDNA BR Quantification Assay Kit (Invitrogen) then pooled with equal DNA mass from each library. The final pooled libraries were submitted to the UCSF Center for Advanced Technology for sequencing on an Illumina Novaseq X 10B sequencer.

### RNA sequencing data analysis

Transcript abundances were quantified based on version ucsf_hc.01_1.G217B of the Histoplasma G217B transcriptome (S5 Data of Gilmore et al^19^). Relative abundances (reported as TPM values^69^) and estimated counts (est_counts) of each transcript in each sample were estimated by alignment free comparison of k-mers between the reads and mRNA sequences using KALLISTO version 0.46.2^70^. Further analysis was restricted to transcripts with estimated counts ≥ 10 in at least three samples. Differentially expressed genes were identified by comparing replicate means for contrasts of interest using LIMMA version 3.46.0^71,72^. Genes were considered significantly differentially expressed if they were statistically significant (at 5% FDR) with an effect size of at least 1.5X (absolute log_2_ fold change ≥ 0.585) for a given contrast.

### Comparative transcriptome analysis

For fungi with RNAseq data available for hyphal and yeast (or spherule, for Coccidioides) (Table S5), reads were downloaded from the SRA. Corresponding transcriptome sequences were downloaded from the sources indicated in Table S5 Data and indexed for KALLISTO. Transcripts were quantified for each sample with KALLISTO version 0.46.2^70^, invoked as:

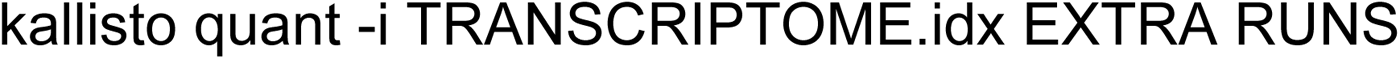

where TRANSCRIPTOME.idx is the appropriate index file, RUNS are gzipped FASTQ files for all runs corresponding to the given sample, and EXTRA are additional flags for appropriate experiment-specific handling of single-ended and strand-specific reads, as given in the “extra flags” column of Table S5. Further analysis was restricted to transcripts with ≥ 10 counts in at least half of the samples from a given dataset. Yeast/hyphal or spherule/hyphal contrasts and BH adjusted p-values were estimated using LIMMA version 3.46.0^71,72^ and genes were considered signficantly differentially expressed if they were statistically significant (at 5% FDR) with an effect size of at least 2x (absolute log_2_ fold change ≥ 1). Histoplasma ortholog groups were taken from S13 Data of Gilmore et. al.^19^. Othologous genes between G217B and the remaining species were determined by InParanoid version 1.35^73^. We used Fisher’s exact test with BH multiple hypothesis correction to test for significant enrichment of hyphal-enriched orthologs from the comparison transcriptomes in the set of 144 HYF1+aneuploid-enriched genes. Kernel density estimates of these distributions are plotted in Fig 7G using the gaussian_kde function from SciPy^74^.

### Generation of increased copy number and overexpression strains

To construct *Histoplasma* strains with increased copy number of specific regions (regions indicated by black bars in Fig. S6A-E, S7D), these regions were amplified from extracted genomic DNA (supplemental table 2). Regions were then cloned into a *Histoplasma* entry vector containing telomere repeats and the gene *URA5*. Transformation of these linearized plasmids into *Histoplasma* was performed as previously described^75^ into WU15-based strains. Strains containing ectopic plasmids with native regulatory elements were used for RNA sequencing shown in Fig. 7. For *HYF1* and *HYF2*, overexpression plasmids were similarly made by amplifying the region indicated in grey in Fig. S6D and S7D.

### Other analysis and statistics

Venn diagrams were made using venn3, except the diagram in 7A which was made using eulerr. Heatmaps were generated in Java TreeView 1.1.6r4^76^. Survival curve was generated using Prism. Throughout this paper, p-values equal to or above 0.05 are considered not significant, p-values < 0.05 are indicated by *, p-values < 0.01 are indicated by **, p-values < 0.001 are indicated by ***. Specific statistical tests are noted in figure legends and methods.

## Data availability

DNA and RNA sequencing reads were submitted to GenBank under BioProjects PRJNA1256922, PRJNA1257453, PRJNA1257851, PRJNA1259587, and GEO series GSE296938. Genome assemblies from BioProjects PRJNA682645 and PRJNA1196486 were used. Previously published sequencing of natural isolates from BioProjects PRJNA416769, PRJNA868688, and PRJNA1003095 were also used for analysis. Accessions for data sets used in figure 7G are given in supplemental table S5. Supplemental table S1 lists strains used in this study.

## Acknowledgements

We thank Daniel Matute, William Goldman, and Victoria Sepulveda for generously sharing clinical *Histoplasma* isolates. We thank Praise Oo for initial growth and assessment of these clinical isolates. We thank Sandy Johnson, Carol Gross, Joe Bondy-Denomy, and Suzanne Noble for useful suggestions. We thank members of the Sil and Noble labs for helpful discussions. The UCSF Center for Advanced Technology and Chan-Zuckerburg Biohub-San Francisco supported this research through use of equipment and/or sequencing.

